# Quantitative analysis of synaptic pathology and neuroinflammation: an initial study in a female rhesus monkey model of the “synaptic” phase of Alzheimer’s disease

**DOI:** 10.1101/251025

**Authors:** Danielle Beckman, Kristine Donis-Cox, Sean Ott, Mary Roberts, Lisa Novik, William G. Janssen, Eliza Bliss-Moreau, Peter H. Rudebeck, Mark G. Baxter, John H. Morrison

**Author notes:** **Corresponding author:** John H. Morrison, California National Primate Research Center, UC Davis, Davis, CA 95626, USA.

## Abstract

**Background:** Soluble oligomers of the Aβ peptide (AβOs) are toxins that target and disrupt synapses. Generation of AβOs has been recently recognized as a probable initiating event in Alzheimer’s disease (AD), leading to cognitive impairment. There is a translational gap in AD studies, with promising drugs developed based on work in rodent models failing in AD patients in clinical trials. Additionally, although women have a two-fold greater lifetime risk of developing AD compared to men, females have not been a focus of preclinical studies. Thus, we sought to develop a model of AβO toxicity in female rhesus monkeys, to take advantage of the more highly differentiated cortical structure in this species as well as the similarities in the endocrine system between rhesus monkeys and humans.

**Methods:** Repeated intracerebroventricular (i.c.v) injections of AβOs were performed in adult female rhesus monkeys. Controls were unoperated aged matched monkeys. High-resolution confocal microscopy and morphometric analysis of Alexa 568 (A568) filled neurons were used to evaluate synaptic, neuronal, and glial markers in the dorsolateral prefrontal cortex (dlPFC) and hippocampus after AβO injections. Cerebrospinal fluid (CSF) and brain tissue were also collected and analyzed for biomarkers of AD pathology, including: phosphorylated Tau protein (pTau), total Tau, Aβ_1-42_, Aβ_1-40_ and TNF-α levels.

**Results:** Here, we report that AβO injection into the lateral ventricle of the brain induces loss of 37% of thin spines in targeted dlPFC neurons, an area highly vulnerable in AD and aging. Further, AβOs associate with the synaptic marker PSD95, inducing loss of more than 60% of local excitatory synapses. AβOs induce a robust neuroinflammatory response in the hippocampus, far from the injection site, with numerous activated ameboid microglia and TNF-α release. Finally, AβOs increased CSF levels of Aβ_1-42_, pTau Ser396 and pTau Ser199, but not Aβ_1-40_ or total Tau.

**Conclusions:** These initial findings from detailed quantitative analysis of effects of AβO administration on synapses in a female nonhuman primate model are a very promising step toward understanding the mechanism of early AD pathogenesis in the primate brain, and may help develop an effective disease-modifying therapy of high relevance to women’s health.

## Introduction

Alzheimer’s disease (AD) is the most common form of dementia in the elderly, with an estimated number of 5.5 million Americans living with the disease in 2017 *(1)*. Despite considerable investment and effort, this prevalence is expected to increase further in the coming decades, following an aging trend of the world population. The pathogenesis of AD has long been linked to the presence of fibrils of amyloid beta (Aβ) protein, which accumulates markedly in AD brains, forming insoluble amyloid plaques. Although plaques play a role in AD pathology, during the past decade much of the focus has turned to biologically active soluble oligomers of the Aβ peptide (AβOs). These oligomers are potent neurotoxins, known to accumulate in AD brains and in animal models of the disease *(2)*. Soluble AβOs, rather than insoluble fibrils or plaques, trigger synapse failure, now regarded by many as the earliest reflection of AD pathogenesis *(2, 3)*.

There are several transgenic AD models for studying the pathogenesis of the disease, but so far, these models fail to capture many aspects of the disease in humans. Transgenic AD mouse models carry mutations that are associated with early-onset familial forms of AD, which account for less than 5% of the cases of the disease. The vast majority of AD cases are sporadic, with a poorly understood etiology that leads to accumulation and impaired clearance of AβO species in the absence of known genetic lesions *(4)*. Another important factor is that most AD studies rely on the use of male animals, although nearly two-thirds of current AD cases are women *(5)* and after the age of 65, the lifetime risk of AD is 1 in 6 for women, whereas it is 1 in 11 for men *(6)*.

Given the role of soluble AβOs in AD and the limitations of currently available animal models, we report here the development of a nonhuman primate model for AD focusing exclusively in females. For this, two adult female rhesus monkeys received repeated i.c.v injection of freshly prepared and fully characterized AβOs, twice a week. Synaptic health, neuron and microglia morphology and CSF biomarkers were evaluated. Our data suggest that exogenous administration of AβOs in female rhesus monkey results in markers of synaptic dysfunction and neuroinflammation that recapitulate features of preclinical Alzheimer’s disease.

## Results

### AβOs induce reduction in thin spine density and morphology in monkey Prefrontal Cortex

Synaptic dysfunction and loss caused by AβO binding to neurons has been recently considered one of the earliest events in the pathologic cascade of AD *(7, 8)*. The dlPFC of primates is highly vulnerable to aging and AD, especially Brodmann area 46, where synapse loss and structural alterations occur in aging with or without neurodegenerative disease *(9–11)*. Analysis of dendritic length of pyramidal neurons in layer III of area 46 in the dlPFC of AβO-injected monkeys showed an increase in the total dendritic length of the neurons, compared to neurons analyzed from age-matched controls (Fig. 1A-B). Although it is not clear if the increased dendritic length happens in basal dendrites, apical or both (Fig. 1C-D), one possibility is that diffusible oligomers damage neurons and induce neurite sprouting as a plastic response, as seen in these same neurons in young ovariectomized female monkeys *(12)*. In addition, increased dendritic length has been observed in a transgenic AD model *(13)* and also in cultured neurons treated with Aβ aggregates *(14)*. Interestingly, increased dendritic trees have also been associated with normal aging as a mechanism to compensate for some local loss of neurons *(15)*.

**Figure 1:**
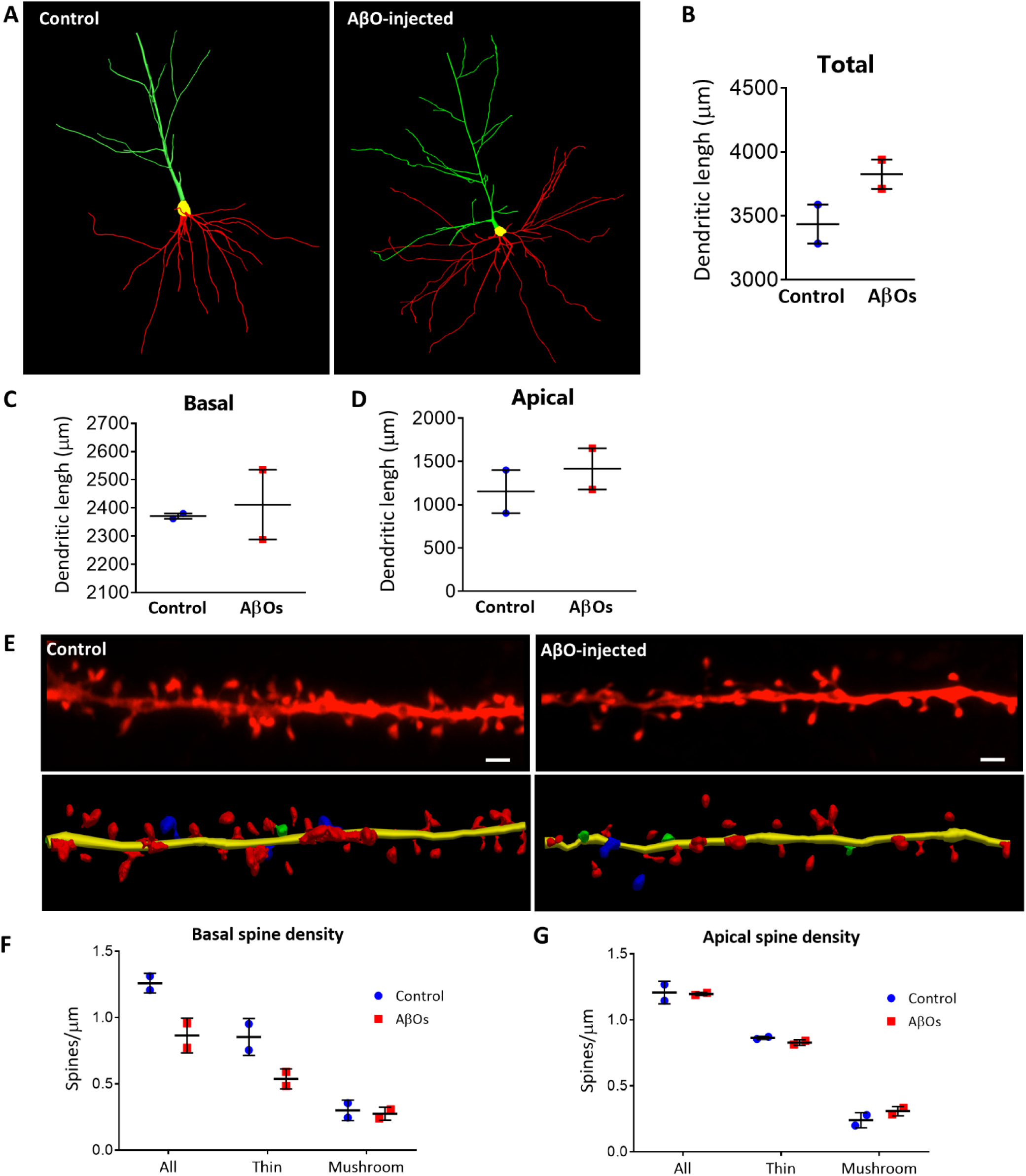
AβO infusions induce morphological changes in the dlPFC pyramidal neurons of rhesus monkeys. (A) Representative 3D neuronal reconstruction of neurons from a control and AβO-injected monkey showing an increased dendritic length in the AβO-injected case. Green: Apical dendrite, Red: Basal dendrites. (B) Increased total dendritic length (μm) in neurons from monkeys exposed to AβOs. Basal (C) and apical (D) total dendritic length comparison between control monkeys and AβO-infused monkeys. (E) Representative projected z-stack and 3D reconstruction of basal dendrite segments showing reduction in the number of spines in a dendritic segment from a dlPFC neuron in the brain of an AβO-infused monkey compared to a control monkey. Red: thin spines, Blue: mushroom spines, Green: others. Scale bar: 2 μm. (F, G) Quantification of the total number of basal (F) and apical (G) spines in dlPFC pyramidal neurons from control and AβO-infused monkeys.

Spine analysis of the same neurons revealed a decrease in the total number of spines, correlating with a decrease in the number of thin spines, but not mushroom spines, in basal, but not apical dendrites (Fig 1D-F). Small thin spines are highly motile and thought to be required for acquiring new information, whereas large mushroom spines are more stable and usually thought to regulate stable memory circuits *(16)*. Our group showed that thin spines of layer III pyramidal neurons are preferentially lost in monkeys with age-related cognitive impairment *(10, 11)*, and can be restored by estradiol treatment*_(12)_ (17, 18)*. Here we show this same population of spines is also vulnerable to toxic AβO species that play a major role in the first stages of biochemical dysfunction in AD.

### AβOs induce loss of PSD95 in pyramidal neurons

Several lines of evidence indicate that deregulation of excitatory transmission can underlie the beginning of synaptic failure in AD. The thin spines are NMDA receptor (NMDAR) dominated *(19)* and vulnerable to the toxic effects of AβOs, which can directly activate this receptors and induce excitotoxicity, leading to neuronal death *(20, 21)*. Postsynaptic density protein 95 (PSD95) is a major scaffold protein that is enriched with NMDAR, and regulates structural and functional integrity of excitatory synapses. We observed that AβOs deposit with high affinity to pyramidal neurons in the dlPFC of monkeys, and lead to a reduction of 63% of PSD95 puncta in the presence of AβO binding. No AβO labeled neurons were observed in age-matched control monkeys, indicating intraneuronal AβOs were not present in the normal healthy adult monkey brain (Fig. 2A-C).

**Figure 2:**
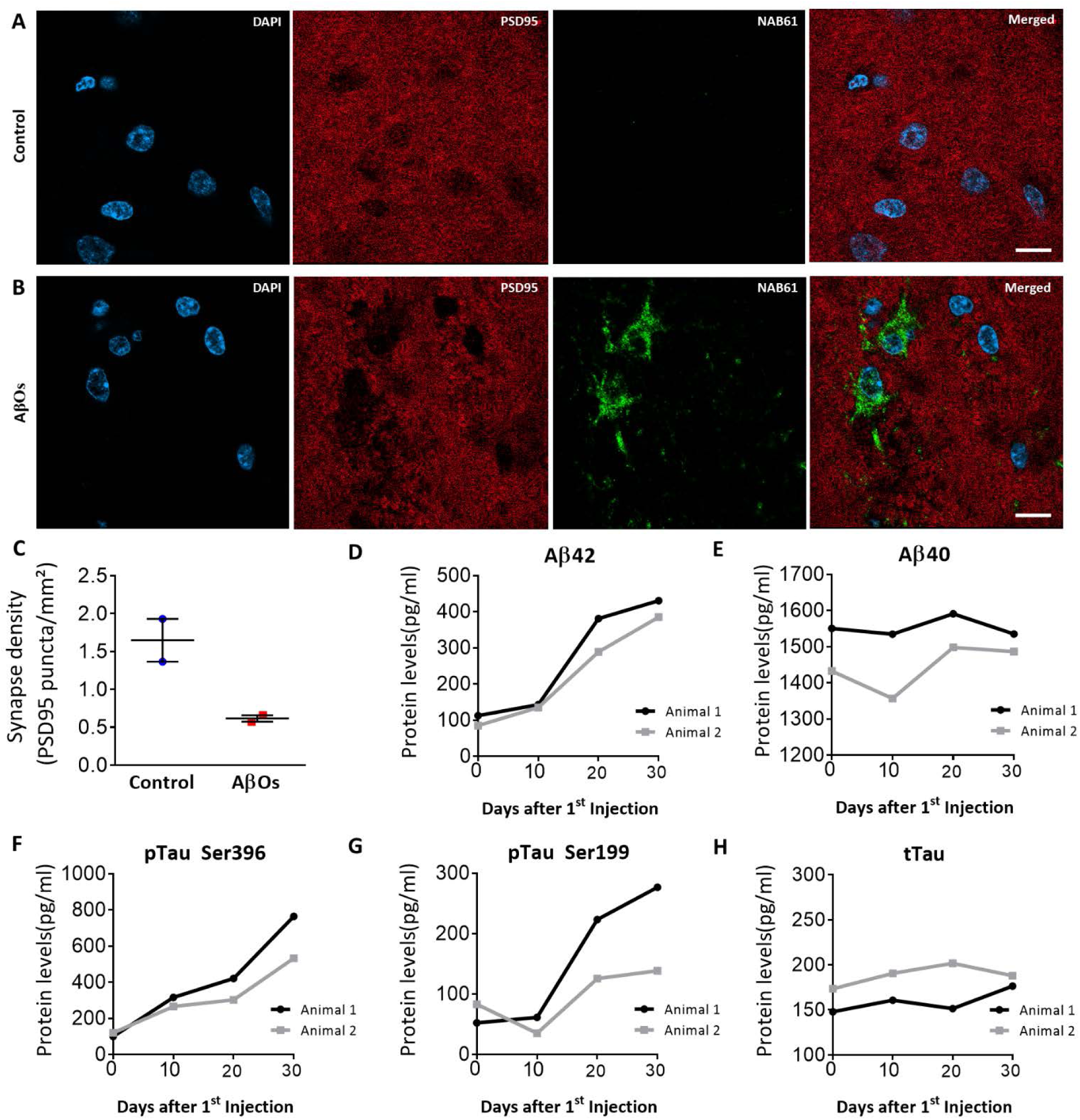
AβOs induces synaptic dysfunction and increase AD biomarkers in the CSF. (A, B) Representative micrographs showing AβOs (NAB61) associate with PSD95 inducing local reduction of the total number of excitatory synapses. Scale bar: 10 μm. (C) Quantification of PSD95 puncta loss in the presence of toxic AβOs. (D-H) AβO-injected female rhesus monkey presents equivalent levels of core biomarkers of AD in humans.

### AβO infusion lead to increase in AD related biomarkers in the CSF

CSF is an optimal source for AD biomarkers since it is in direct contact with the extracellular space of the brain, and can directly reflect biochemical changes in the living brain. Core biomarkers such as Aβ and tau levels can be used to identify the progress of the pathogenic events in AD as early as 25 years before expected symptoms *(2, 22)*. Because of this, CSF was collected every ten days during the AβO injection protocol. Aβ_1-42_, but not Aβ_1-40_ was increased (pg/ml) during injections in AβO treated monkeys (Fig 2D, 2E). Interestingly, tau phosphorylation at two sites: serine 396 and serine 199 also increased during the period of injections, but total Tau protein levels did not change (Fig 2F-H). Even though pTau was increased in the CSF, no neurofibrillary tangle formation was detected in the brain. This contrasts with a previous report from Forny-Germano et al, in cynomolgus monkeys injected with AβOs into the cerebral ventricles *(23)*. Importantly, all the core biomarkers evaluated in the rhesus monkey CSF presented equivalent values compared to human baseline CSF levels or proteins *(22, 24)*.

### AβOs trigger neuroinflammation in the hippocampus

Recently, preclinical data have shown that glial cell activation and neuroinflammation accompanies AD pathology and contributes to the development of the disease. Microglia, the resident immune cells of the brain, when activated by the presence of Aβ species can be harmful to neurons, mediating synapse loss by engulfment of synapses, activating neurotoxic astrocytes and secreting inflammatory cytokines *(25–27)*. This process can occur in different types of neurodegenerative disease, but is especially relevant for AD pathogenesis since the majority of risk genes for AD are highly expressed in microglia. To investigate microglia morphology and function in response to AβO injection, we analyzed the dentate gyrus (DG) of the hippocampus of AβO-injected monkeys and controls, an area far from the injection site, thereby avoiding any inflammatory process caused by the injection needle penetrating the cortex over the lateral ventricle. The DG of the AβO-injected animals presented a robust microglial amoeboid population, as determined by staining with microglial IBA-1 marker (Fig. 3A, 3B).

**Figure 3:**
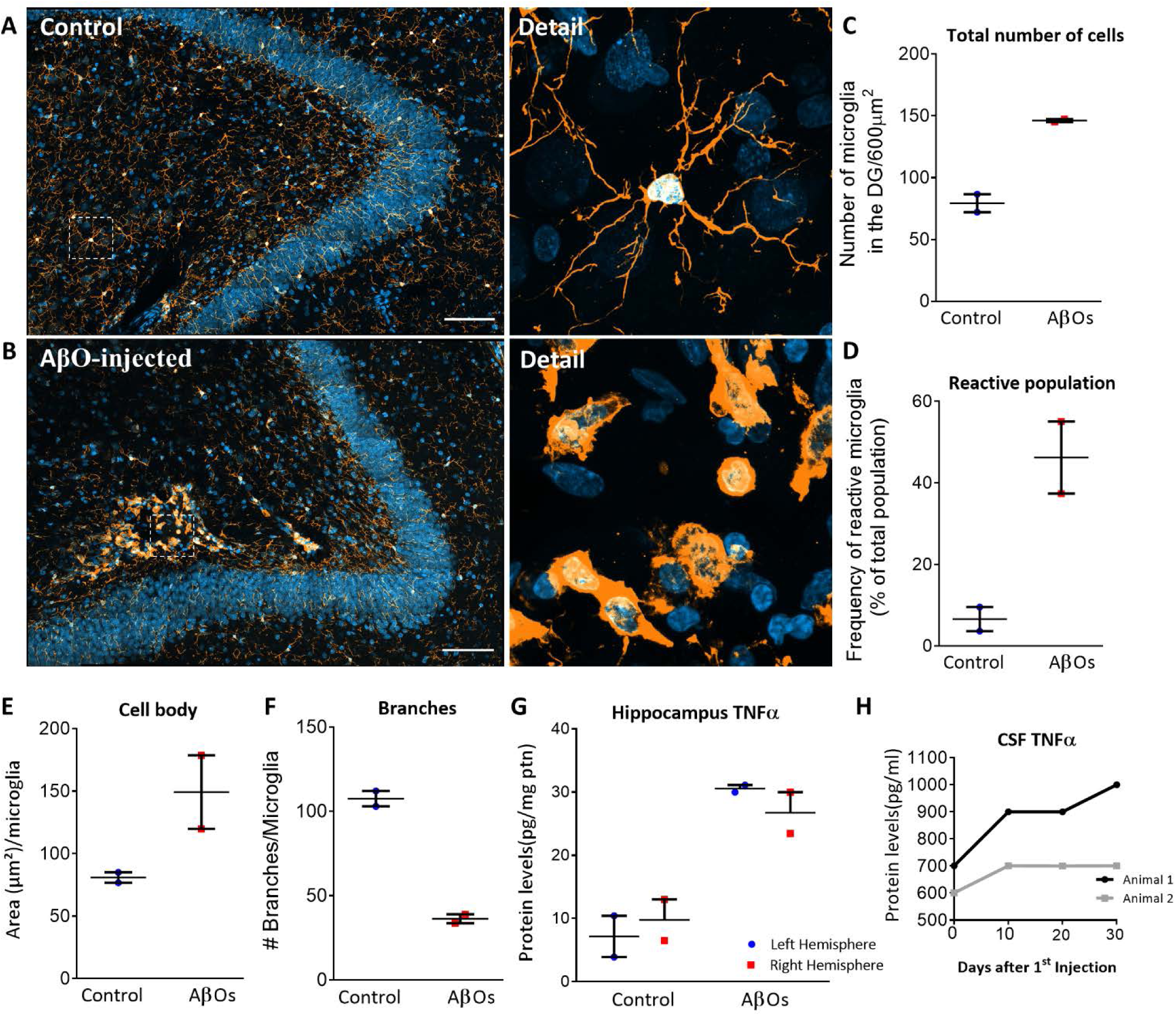
AβO injection induces robust microgliosis and neuroinflammation in the hippocampus. (A,B) Representative z-stacks staining with IBA-1 for microglia showing profound morphologic changes and microgliosis induced by AβOs in the CA3/hilus of the hippocampus. Scale bar: 100 μm. (C) Total number of microglial cells is increased in AβO-injected animals in comparison with controls. (D) Reactive microglial population is increased more than 7x in AβO-infused monkeys. AβOs also increased cell body size (E) and decreased the number of branches (F), indicative of intense microgliosis. (G) There was more than a threefold increase in the proinflammatory cytokine TNF-α in the hemispheres of AβO-injected monkeys. (H) TNF-α levels in the CSF were increased in one of the two AβO-injected monkeys.

Usually healthy microglia present extended and ramified processes, but when activated, there is an increase in the cell body and branch retraction. We observed an increase of more than 80% of the total number of microglia cells in the DG of AβO-injected monkeys in comparison to age-matched controls, suggesting cells are being recruited from other brain sites (Fig. 3C). The amoeboid (reactive) population constituted almost half (46%) of the total number of microglial cells in the DG in AβO-injected monkeys, in contrast with 7% of the total population in control monkeys (Fig. 3D). Activated microglial cells in AβO-injected monkeys also presented morphological changes, with an 84% increase of cell body size (Fig 3E) and 34% decrease of total branching compared to the same area in control monkeys (Fig 3F). Finally, hippocampal samples analyzed for cytokines showed a prominent increase in TNF-α levels when compared to control animals (Fig 3G). Interestingly, no difference was observed between left and right hemispheres of all animals, suggesting AβO diffuses through the whole brain and inflammatory reactions are not constrained to the hemisphere where the lateral ventricle infusions occur. TNF-α levels were below the limit of detection in other brain areas (data not shown) and markedly increased only in the hippocampus. Finally, TNF-α levels were increased in the CSF in one AβO-injected monkey, suggesting a localized neuroinflammation in the hippocampus can constitute an early event in AD pathology.

## Discussion

Here we report a quantitative analysis of synaptic dysfunction and neuroinflammation in an initial study developing a female nonhuman primate model of AD. Using several sophisticated microscopy techniques, including 3D reconstruction and quantitative analysis of A568-loaded cells in conjunction with high resolution confocal microscopy, we provide an extensive and detailed analysis of neuronal and microglial morphology in the dlPFC and hippocampus of AβO-infused monkeys. We demonstrated here that when injected in the lateral ventricle, AβOs diffuse and accumulate in areas of the monkey brain relevant to higher-order cognition, including the dlPFC and hippocampus, critical areas in AD pathology *(28)*. Neurons in the dlPFC, vulnerable to aging and neurodegenerative diseases *(29)*, showed increased dendritic branching and specific loss of thin dendritic spines. These effects are more pronounced forms of the synaptic disruptions we have observed in cognitively impaired aged monkeys without exogenous AβO infusions *(29)*, and are suggestive of early synaptic dysfunction in AD.

AβO infusions also induced reactive microgliosis in the hippocampus, far from the injection site, with localized increases in levels of the proinflammatory cytokine TNF-α. In pathological conditions, microglia release large amounts of TNF-α, which increases inflammatory processes, synaptic pathology, and excitotoxicity *(30, 31)*. Increased TNF-α levels were found in the CSF of one of the AβO-injected animals, but not the other, possibly due to the short period of injections used in this study.

It is known that in humans and in nonhuman primates, the loss of circulating estrogen occurring normally with menopause exacerbates normal age-related deficits *(18, 32, 33)*. Here we show that infusion of AβOs in adult female rhesus monkeys can induce alterations reminiscent of early stages of AD pathology, including synaptic dysfunction and microgliosis. This is especially relevant because accumulation and oligomerization of Aβ_1-42_ usually happens decades before manifestations of cognitive and behavioral impairment. Thus, we provided here a reliable platform to develop future interventions focusing on counteracting initial pathogenic effects in AD, and improving women’s health.

## Conclusion

Our initial data indicate that exogenous administration of AβOs into the lateral ventricle of adult rhesus monkeys results in markers of synaptic dysfunction that recapitulate many of the hypothesized features of preclinical Alzheimer’s disease. Such synaptic data cannot be obtained from humans, but tight linkage can be obtained in an AD nonhuman primate model. Following this step, the potential role of ovarian hormone replacement in aged females will be evaluated, as this is one major concern in women undergoing hormonal changes during menopausal transition. Given the translational power of the female AD monkey model, such data will have direct relevance to women’s health and AD, and provide a novel model for testing new interventions.

### Authors’ contributions

DB, KDC, SO, MR, LN, WJ, PHR, and MGB performed experiments. DB and JHM analyzed the data. JHM, MGB and EBM designed the research. DB, MGB and JHM wrote the manuscript.

## Acknowledgements

We thank Silvia Hilt and John Voss for the guidance with AβO characterization, Anne Gibbons for the help with perfusions, and Virginia Lee for the kind gift of Aβ oligomer-specific NAB61 antibody. We also thank Sarah Motley for the guidance with data analysis, Amanda Dao, Marc Friedman and Eric Zhou for the help with neuronal analysis and Jeffrey Roberts for insightful discussions. The authors have no conflict of interest to disclose. This project was supported by National Institute on Aging (NIA) award P01-AG016765 and by a pilot grant supported by National Institutes of Health (NIH) Office of the Director award P51-OD011107. The California National Primate Research Center is supported by NIH Office of the Director award P51-OD011107. The content is solely the responsibility of the authors and does not necessarily represent the official views of the NIH.

## Material and methods

### Animals

All experiments were conducted in compliance with the National Institutes of Health Guidelines for the Care and Use of Experimental Animals approved by the Institutional Animal Care and Use Committee at the University of California–Davis (Protocol number 18994). Four female adult (age range, 11–19 years old) rhesus monkeys (*Macaca mulatta*) were used for morphometric analyses and biochemical assays.

### Surgery

Under full aseptic conditions and in a dedicated operating suite, two monkeys were implanted with a plastic chamber (Crist Instruments, MD) that allowed access to the dural surface. Monkeys were premedicated with an IM injection of ketamine (10 mg/kg) and dexmedetomidine (10 μg/kg). The monkeys were intubated, placed in a stereotaxic frame, and maintained on isoflurane anesthesia (1.5%, to effect). Intraoperative monitoring included pulse oximetry, capnography, EKG, and blood pressure. The head was scrubbed and draped for aseptic surgery. To expose the skull over the midline, the skin and underlying galea were incised and the muscles retracted. A craniotomy was made corresponding in size to the inner diameter of an MRI-compatible plastic infusion cylinder (Crist Instruments), centered 20-25 mm anterior to the interaural plane. The cylinder was fixed to the skull with titanium screws (Venterinary Orthopedic Implants) with a small amount of dental acrylic (Orthojet), and the galea and skin were then closed around the implant. Postoperative analgesia consisted of oxymorphone.

### MRI

Post-implant MRI scans were conducted to obtain precise coordinates for placement of injections into the lateral ventricles. At the time of the scan, a grid was placed into the chamber filled with dilute gadolinium. Dilute gadolinium is a contrast agent (0.01 – 1%) that facilitates visualization of the grid within the implant. Infusion needles and cannula were custom made for each animal using the measurements determined from coordinates obtained from the MRI.

### Infusions

Animals were premedicated with ketamine and dexmedetomidine. The animals were placed on flow nasal oxygen and vitals were monitored. The chambers were first cleaned with 500 mls of sterile saline plus dilute betadine solution or dilute chlorhexidine solution using sterile technique. This was followed by a final chamber rinse of 12 mls of pure sterile saline. Next the sterile grid was placed into the chamber and secured in place being careful to use the exact orientation from the MRI scan. The sterile cannula was carefully lowered through the grid and into the brain using the coordinates determined from the MRI. The AβO or control was drawn up into a sterile tuberculin syringe. The syringe was attached to the sterile infusion needle assembly. The line between the needle and the syringe hub was allowed to fill with the infusion material from the syringe until a tiny bead was observed at the tip of the syringe. This was done to avoid injecting air into the ventricles. The needle was carefully lowered into the brain through the cannula. The infusion material was delivered at a rate of 25 microliters every 30 seconds. The infusion was followed with less than 100 microliters of sterile saline to clear the line going from the syringe to the needle to ensure that the total amount had been delivered into the ventricle. After waiting 2 minutes the needle and the cannula were removed from the brain. When infusion was complete a clean sterile cap was placed on the chamber. Lastly, the animals were reversed with a dose of atipamezole and monitored during recovery.

### AβO preparation

Synthetic Aβ_1-42_ human peptide (California Peptide) was used to prepare Aβ solutions as previously described *(34)*. Briefly, Aβ films were prepared dissolving 1 mg of peptide in 270 μL of ice cold HFIP. Tubes were allowed to evaporate in a laminar flow hood overnight, free of visible particles. Films were stored at −20°C and used within 10 days after preparation. AβOs solutions were prepared dissolving films in 10 μL of DMSO (5mM) and 490 μL of cold PBS. The solutions were vortexed and incubated at 4°C for 24 hours to allow oligomerization. Solutions were freshly prepared every day immediately prior injections. Protein levels were determined to verify allow a mass of 100 μg of AβOs or scramble Aβ per injection.

### AβO characterization by dynamic light scattering

Particle size analysis of the Aβ oligomers solutions were carried out for every preparation prior to injections using dynamic light scattering (DLS) with a Brookhaven 90Plus instrument that monitors scattered light at 90° to the excitation. The samples were centrifuged at 14000xg to remove the large amorphous aggregates just prior to measurement. Measurements were made at sequential dilutions from the undiluted sample of 5μM to 1:2, 1:4, 1:10, 1:100 dilution factors. The average hydrodynamic radius was calculated using the ZetaPlus Particle Sizing Software Version 3.57 (Brookhaven Instruments).

### Perfusion and tissue preparation

All rhesus monkeys were deeply anesthetized with ketamine hydrochloride (25 mg/kg) and pentobarbital (20–35 mg/kg, i.v.), intubated, and mechanically ventilated. The chest was opened to expose the heart and heparin was injected into the left ventricle to facilitate vasodilation. The descending aorta was clamped immediately following intubation, and the monkeys were perfused transcardially with warm 0.9 % saline for 5 min followed by cold 0.9% saline for approximately 5-7 min at a rate of 220 ml/min. After perfusion, the brain was removed and completely submerged in ice cold saline for 10 min. After this period, the brain was placed into a monkey brain mold and coronal slabs were prepared for regions of interest with both fixed and frozen blocks prepared for morphological and biochemical studies, respectively. Samples for microscopy were immersed in fixative (4% paraformaldehyde in phosphate-buffered saline, 7.4 pH + 0.125% glutaraldehyde) for 36 hours before being rinsed in buffer and processed. Frontal and temporal blocks were cut serially on a vibratome (Leica) in 400 μm-thick sections (for intracellular injection of A568; Invitrogen) or 50 μm-thick sections for fluorescence confocal microscopy. For biochemical analysis, 400 μm-thick sections were sonicated and diluted using proteases and phosphatases inhibitors an immediately stored at −80°C until analysis.

### Immunohistochemistry and microscopy image analysis

#### Quantitative analyses of spine density and spine morphology

Intracellular injection of layer III pyramidal cells with A568 and quantitative analysis was performed according to methods previously described *(10, 12)*. Dendritic segments were chosen in a systematic random fashion for branching and spine analysis. A total of 20 apical and 20 basal segments were acquired for each experimental condition, with a total of 6-10 neurons being analyzed per group. Photomicrographs of the neurons and segments were made using a Zeiss LSM 800 microscope with Airyscan. After acquisition, neurons and segments were exported to Neurolucida 360 (MBF Bioscience) for 3D reconstruction and spine analysis. Data was then imported into Matlab for spines classification into thin and mushroom types based on categories previously established in our lab *(10)*.

#### PSD95 and AβOs puncta

50 μm-thick free-floating sections were incubated overnight with monoclonal oligomer-specific NAB61 (provided by Virginia Lee, 1:2000), PSD-95 (Cell Signaling, D27E11, 1:1000) and NeuN (Abcam, ab134014) antibodies. Tissue were then thoroughly washed with PBS and incubated with Alexa Fluor secondary antibodies (Invitrogen, 1:500) for 2 h, at room temperature. Slides were mounted with Prolong Diamond Antifade with Dapi (Invitrogen). For each experiment: 20 single images were acquired from each independent experimental condition, in a blinded fashion at 63x using a Zeiss LSM 800 microscope. Staining was analyzed using Puncta Analyzer v2.0 plugin (ImageJ software). Neurons were identified using NeuN staining and PSD95 puncta was analyzed in the area around the neuron that presented positive NAB61 staining. For controls, a total area around imaged neurons was selected identical to experimental neurons for PSD95 puncta quantification.

#### Microglia morphology and quantification

For microglial analysis, 50 μm-thick free-floating sections were incubated overnight with IBA-1 (WAKO, #019-19741, 1:1000) antibody. Immunohistochemistry was performed as described above. Three composite confocal images consisting of 70 tile at 25 μm z-stacks thickness were imaged at 63x to capture the entire DG using the Zeiss LSM 800 microscope for each animal using different portions of the hippocampus. The maximum intensity projection of the IBA-1 staining was converted to a binary image. Only cells presenting the entire cell body were isolated followed by morphometric analysis using the AnalyzeSkeleton plugin (ImageJ software).

### ELISAs

Tissue samples were homogenized in a buffer containing 100 mM Tris, 150 mM NaCl, 1 mM EGTA, 1 mM EDTA, and 1 % Triton X-100, supplemented with protease and phosphatase inhibitor cocktails. Aβ_1-42_, Aβ_1-40_, tTau, pTau Ser 396, pTau Ser199 (Invitrogen) and TNF-α (Millipore-Luminex) ELISAS assays were performed according to each kit manufacturer instructions, following sample dilution optimization.

